# Identification of ketotic cattle based on milk spectroscopic characteristics: a chemometric modelling approach built upon the principles of Aquaphotomics

**DOI:** 10.1101/2024.12.18.629140

**Authors:** Simone Giovinazzo, Carlo Bisaglia, Tiziana M.P. Cattaneo, Laura Marinoni, Giovanni Cabassi, Massimo Brambilla

## Abstract

The early diagnosis of metabolic disorders in intensive farms, such as ketosis, allows a timely health treatment, resulting in improved animal welfare and reduced productivity and economic losses for farmers. This approach represents an opportunity for more sustainable animal production practices.

For these reasons, we developed a chemometric processing model based on GNU Octave capable of identifying milk sample categories based on the chemo-physical characteristics of their near-infrared signature. Such a characterisation relies on the innovative approach of aquaphotomics, which investigates the interaction between light and water molecules in any biological system. Aquaphotomics leverages the properties of water as a biomarker and amplifier to identify changes in the system indicative of a perturbation, such as the onset of a metabolic disorder.

The realised package allows spectral data elaborations, including preprocessing methods like the calculation of spectrum derivatives, noise removal and scatter correction techniques, as well as chemometric analysis like Principal Component Analysis (PCA), Linear Discriminant Analysis (LDA), Quadratic Discriminant Analysis (QDA), and Partial Least Squares Regression (PLSR). In this way, the code enables the identification of the most significant water bands implicated in the origin of the disease, with the potential to represent an innovative approach for monitoring the health status of animals.

## Introduction

Aquaphotomics is a scientific discipline that combines principles of spectroscopy, chemometrics, and water science to monitor the properties and functionality of any system (Van De Kraats et al. 2019). Water molecules constitute a dynamic network of hydrogen bonds interacting with any solute dissolved in solution. This framework is susceptible to any changes in the surrounding environment, thus acting as a molecular mirror that systematically detects and interprets the result of a perturbation. Such alterations result in specific changes in the molecular conformations of water molecules clusters, which are reflected in measurable spectral variations in the near-infrared (NIR) absorbance pattern of water (Kovacs et al. 2022). Therefore, in aquaphotomics, the spectral model of water is used as a sensor that reflects the cumulative effect of all the system components, allowing an in-depth and integrated analysis of the behaviour and interactions in the studied system (Muncan & Tsenkova, 2019).

Aquaphotomics could constitute a promising method for continuous and real-time monitoring of the health status of farmed animals, as demonstrated by veterinary diagnostic studies conducted for the early identification of mastitis in dairy cattle (Tsenkova & Muncan, 2021). In particular, the water absorption bands in the NIR region of the spectrum can serve as biomarkers to detect subtle biochemical changes associated with the initial stages of a disease. This approach could represent a reliable tool to promote dairy cattle’s health and physiological welfare through sustainable agricultural practices as, for example, in the early diagnosis of ketosis.

Ketosis is a metabolic disorder characterised by elevated levels of ketone bodies (mainly beta-hydroxybutyrate and acetone) in the blood, resulting from a negative energy balance in dairy cattle (Deniz et al. 2020). This condition can also manifest in a subclinical and silent form, with an estimated incidence that ranges from 20 to 30% (Loiklung et al. 2022). A significant consequence of ketosis is that the mobilisation of ketone bodies negatively impacts production yields, compromising the overall qualitative characteristics of the milk produced: i) the lipid content increases, in particular the concentration of long-chain fatty acids and unsaturated fats; ii) the concentration of milk proteins decreases, including casein; iii) ketosis affects the mineral content, resulting in a decrease in calcium, magnesium and phosphorus, and an increase in sodium and potassium levels (Setti, 2017). Therefore, the overall effect of these variations in biochemical analytes can affect the behaviour of the water molecules surrounding milk analytes, thus offering, using aquaphotomics, the possibility for early diagnosis of conditions, such as ketosis, before the clinical manifestation of symptoms.

Spectral data processing requires the execution of a multivariate spectral analysis and the application of chemometric models to extract the spectral pattern of water absorption caused by the onset of ketosis and to discriminate the physiological state of the animal starting from the absorbance values of the analysed milk samples. The extraction of hidden information from raw spectra includes preprocessing methods, such as the calculation of spectrum derivatives, noise removal and scatter correction techniques; additionally, chemometric analysis plays a crucial role, employing exploratory techniques to reveal patterns and structures in the data, classification methods to group samples according to their spectral characteristics, and regression analysis to link sample spectra to quantifiable properties (Tsenkova et al. 2018). This work aims to present a code developed using the GNU Octave software language program (version 8.4.0 https://www.gnu.org/software/octave/) to classify milk samples according to their near-infrared spectral profiles. A case study is presented here, starting from the developed experimental model based on a real scenario. However, this guideline provides for the analysis of near-infrared spectral data can be extended to any context of analysis and research using the developed code as a reference framework.

## Methodology

### 1. Experimental model

In this experimental study, the code was designed to process spectroscopic data from milk samples collected from three commercial cattle dairy farms. For each sample, compositional parameters (fat, proteins, lactose) were obtained in real-time through the AfiLabTM device (S.A.E. Afikim, Afikim, Israel) (Kaniyamattam & De Vries, 2014), an analyser of milk components based on NIR light refraction. The lactation number of each animal was subsequently obtained from the breeder. Milk samples were initially classified into two groups based on the calculated ratio between fats and proteins. Specifically, samples with a ratio greater than 1.4 were classified in the “Ketotic” group, while those with a ratio lower than the established diagnostic threshold were classified as “Not-Ketotic”. Overall, a total of 207 milk samples were collected, and subsequently analysed, using a portable NIR spectrophotometer (MicroNIRTM OnSiteW - VIAVI Srl, Italy) in transflectance mode. Each recorded spectrum resulted from the averaging of 50 successive scans. The spectral range covered was between 908 to 1676 nm with a resolution of 12.5 nm. Each milk sample was analysed in triplicate, using pure water samples (at room temperature and heated to 40°C) as environmental controls. Table 1 presents the total number of spectra obtained from each farm, categorised into the two metabolic groups.

**Table 1.** Total number of spectra obtained from each farm, classified as “Ketotic NIR spectra” and “Not Ketotic NIR spectra”.

| Name of the Farm | Ketotic NIR Spectra | Not Ketotic NIR Spectra | Total |
| --- | --- | --- | --- |
| Farm 1 (DB) | 42 | 396 | 438 |
| Farm 2 (BR) | 12 | 123 | 135 |
| Farm 3 (NIS) | 0 | 48 | 48 |
| **Total** | **54** | **567** | **621** |

This initial sampling phase resulted in an Excel database, where each row corresponds to a NIR spectrum. The columns include, in order, the values of fat, protein, lactose, the fat-protein ratio, the lactation number of the animal, and the absorbance values recorded for each of the 125 wavelengths analysed (Figure 1). All the samples initially classified as “Ketotic” were placed in the first rows of the dataset, followed by the milk samples belonging to animals identified as “Not-Ketotic”. This ordering was based on the fat/protein ratio, with higher ratios corresponding to ketotic samples.

**Figure 1.**
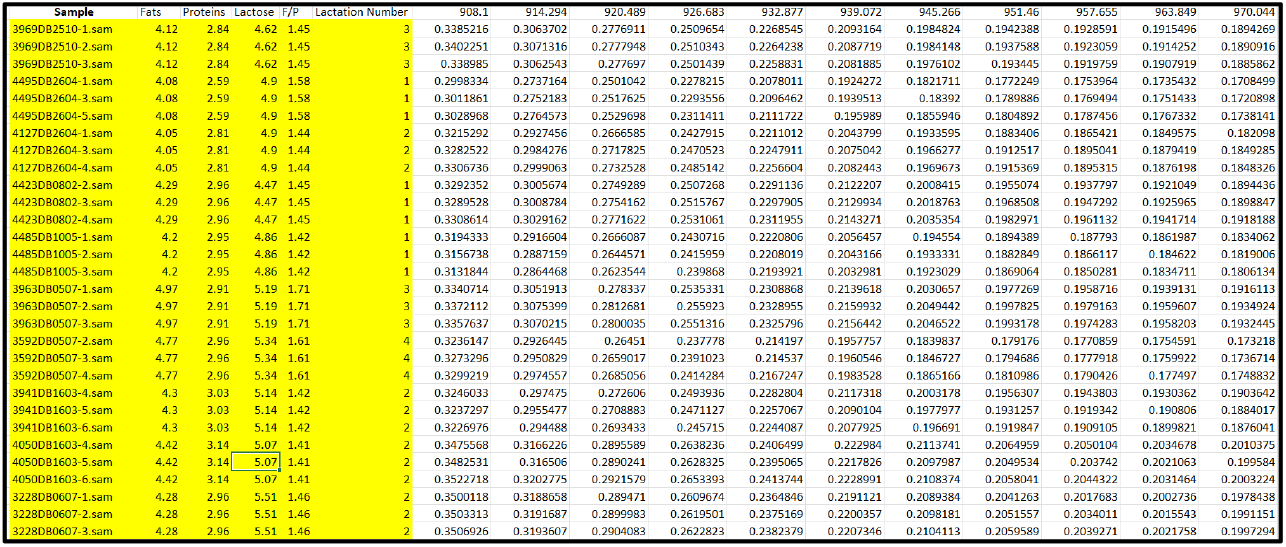
Subset of the collected data obtained from the Excel database.

### 2. GNU Octave Package

GNU Octave is an open-source programming language compatible with the MATLAB syntax and primarily designed for numerical calculations and scientific data analysis. This program was selected to develop an automated processing code for spectroscopic data aimed, in this context, at categorising milk samples according to their spectral profile. Several reasons guided the selection of this platform: i) GNU Octave provides an advanced working environment for matrix calculation and algorithm processing, allowing the management of large datasets and reducing the possibility of error, with considerable time-saving in the execution of complex and repetitive calculations; ii) it supports the import of data in text files format (.txt), as shown in this study, and the export of results in Excel file format (.xlsx); iii) GNU Octave also allows the interactive visualisation of data through plots, offering functions for the overview, zoom and rotation of the graphs, thus facilitating detailed and dynamic analysis of the results.

### 3. The Data Processing Workflow

The data processing workflow comprises two sections, corresponding to distinctive typologies of operations that are summarised in Figure 2:

**Figure 2.**
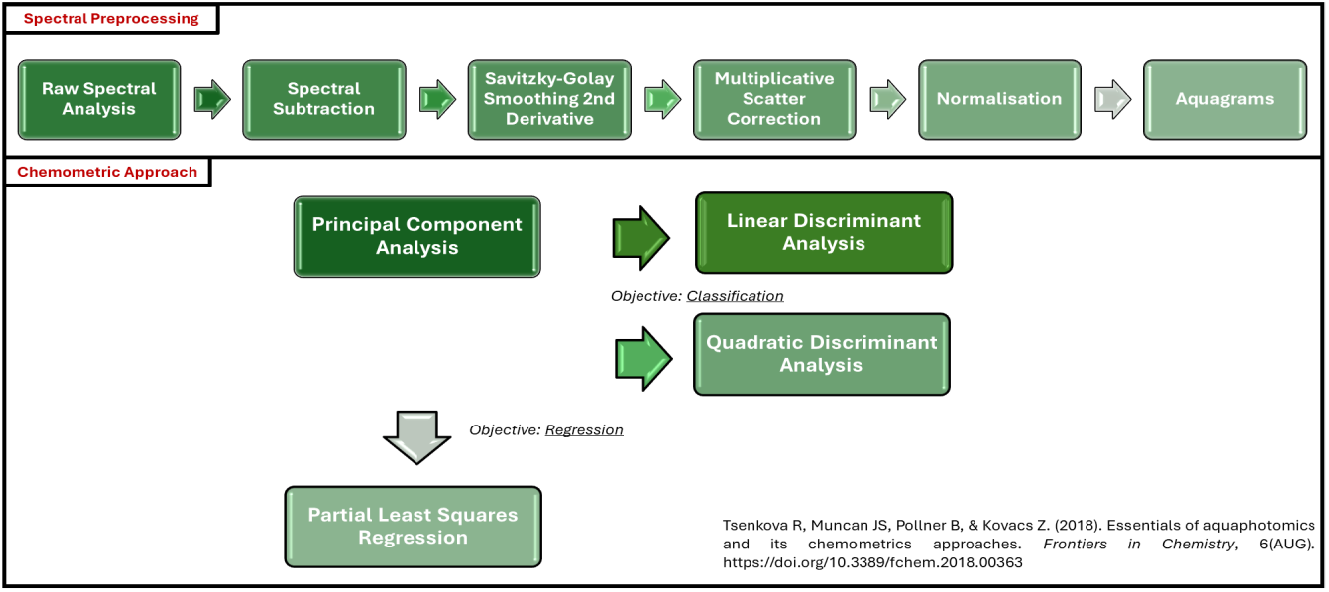
Data processing workflow adopted for milk samples spectral data processing (Tsenkova et al. 2018).

- Spectral Preprocessing, a set of data manipulation operations aimed at improving spectra resolution and noise correction. Measured spectra are often subject to unwanted influences that could compromise measurement accuracy; these may include instrumental noise, environmental interference due to temperature or humidity variations and operator errors (Roger et al. 2022).
- The Chemometric Approach, a multivariate technique utilised to extract significant information from data, especially when there are many variables. This approach includes exploratory analysis of the data to reduce its dimensionality, facilitating the interpretation of relationships and trends (PCA), data grouping techniques to classify elements based on their distance from class centroids (LDA and QDA), and also data modelling methods to develop quantitative models that link measured responses to a set of predictors (PLSR) (Todeschini, 1998).

## Analysis of the Results and Discussion of the Code

The Ketosis_Project code is available from GitLab, using the following link: https://gitlab.com/octave_keto/ketosis_project

This discussion section presents an exhaustive overview of the spectral data operations realised in the code, focusing both on the graphs resulting from the analysis and also on the interpretation of the results. Moreover, in the Supplementary Table S1 it is possible to find a complete list of all the functions utilised in the script, together with a brief introduction of their purpose.

The original Excel database was firstly converted into two text files imported in the code in two distinctive steps:

- *KetosisData*.*txt* is a file containing the spectral data obtained from the NIR instrument, organised as a 665×125 matrix. The first line reports the detected wavelengths, from 908 to 1676nm, while the subsequent lines contain the absorbance values read for each wavelength of the analysed samples; in this specific example, the first 43 lines correspond to the spectra of water, used as an environmental control (19 for room temperature water and 24 water heated to 40°C), and the following 621 lines belong to milk samples spectra.
- *Composition*.*txt* is a file that contains the chemical parameters referring exclusively to the milk samples, organised in a 621x5 matrix. In particular, it includes the fat, protein, lactose, fat-protein ratio values and the lactation number of the analysed cow. The order of samples in both text files is kept unchanged to ensure the accuracy of the joint analyses between spectral data and chemical parameters.

First of all it is fundamental to realise the cleaning of the workspace, carried out with the aim of deleting any previously created variables that could interfere with the script’s performance. It follows the loading of packages to add extra features to the main programming language. In this case, such specialised libraries are necessary for signal processing, statistical analysis and data input/output operations. The actual code starts with the loading of the data contained in ‘KetosisData.txt’ into the “Data” matrix, followed by the assignment of the first row of the matrix (wavelengths) to ‘Lambda’, and the rest of the rows (absorbance values) to ‘Spettri’. Thereafter, the **plot()** function (Figure 3, on the left) creates a two-dimensional graph with x and y axes titled “Raw Spectra” (Figure 3, on the right), which is used to check the raw spectral data of the analysed samples, to verify the existence of prominent absorption peaks at 1190 and 1450 nm as reported by Luck (1973).

**Figure 3.**
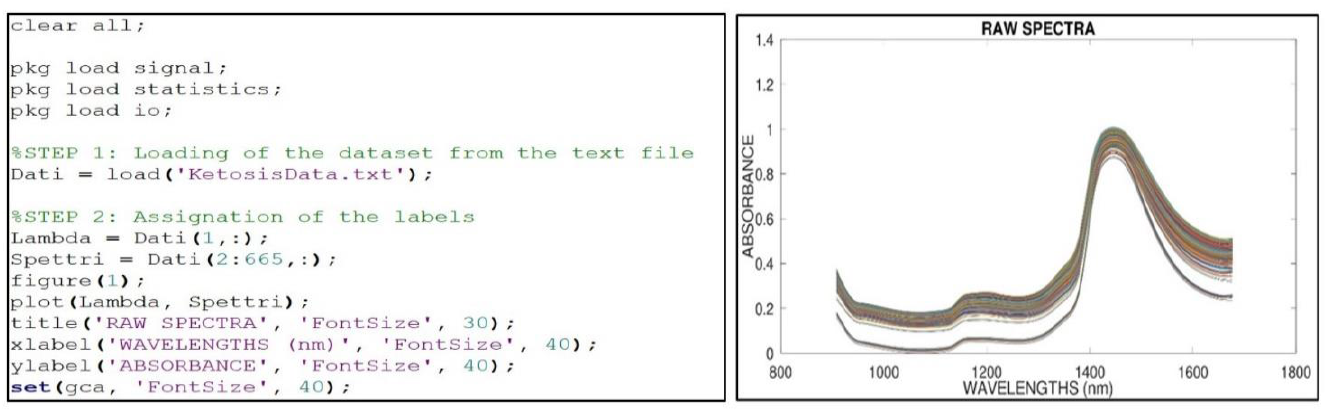
Section of the code used for data processing with the resulting raw spectral graph. On the x-axis the wavelengths from 908 to 1676 nm, while the y-axis reports the experimentally measured absorbance for each sample. Water samples, used as an environmental control, exhibit lower absorbance values and tend to separate from milk samples.

### 1. Spectral Preprocessing

The first preprocessing phase is represented by the spectral subtraction of the mean spectrum of pure room temperature water (approximately 20–25°C) from the analysed milk samples. Room temperature water was chosen as a reference to minimise the spectral contribution associated with temperature, which influences the three-dimensional structure and dynamics of the water molecular network. This phase helps to highlight spectral variations and improve the identification of water bands hidden under broad overtone and combination peaks (Tsenkova et al. 2018). The spectra thus treated are subjected to a subsequent process of smoothing and spectral derivation through the use of the Savitzky-Golay filter. This passage is fundamental both to reduce the influence of noise and to improve the resolution of spectral characteristics, which allows separating, for example, overlapping peaks. These categories of smoothing and derivative operations leverage the GNU Octave function **sgolayfilt()**, a function responsible for applying a 21-point 2nd-order polynomial to the spectral data set (Figure 4).

**Figure 4.**
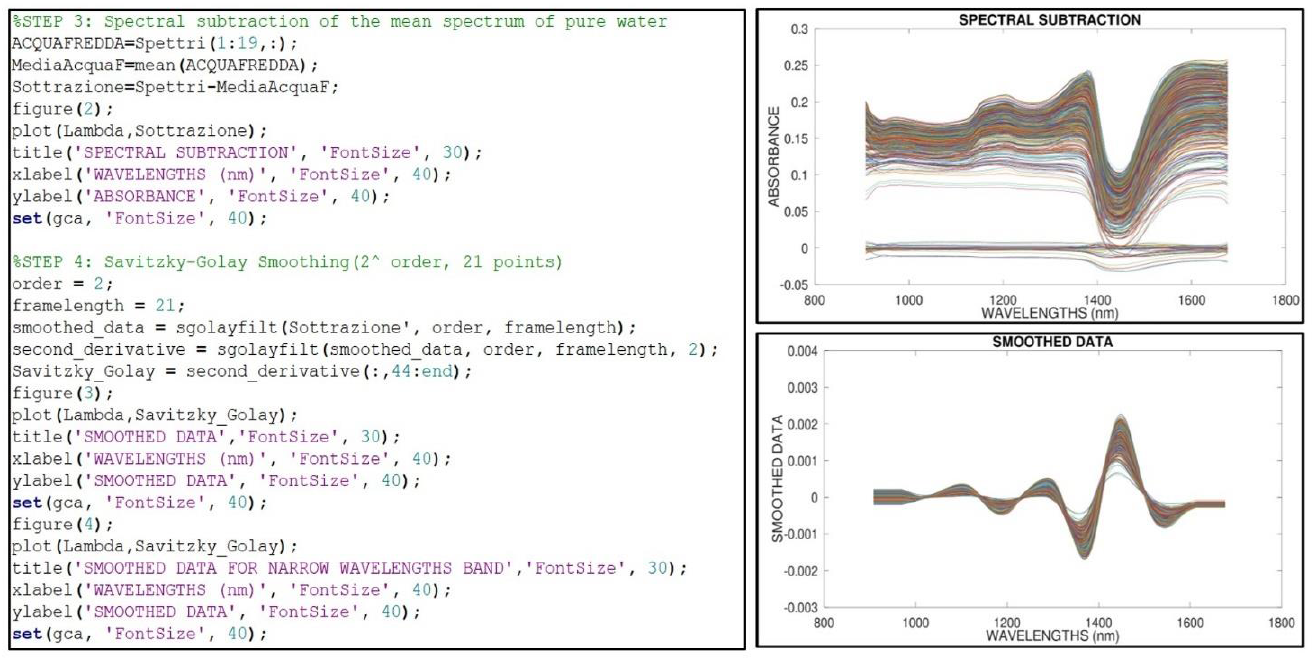
Section of the code used for the first steps of spectral preprocessing. On the top right the graph resulting from the spectral subtraction process, while on the bottom right the graph resulting from the smoothing and derivative calculation.

Preprocessing proceeds with two essential sets of operations: the Multiplicative Scatter Correction (MSC) and the Normalisation. These operations are performed separately for samples categorised as “Ketotic” and “Not-Ketotic” to better highlight the spectral differences between the two groups. First of all, the dataset is reorganised into a three-dimensional matrix thanks to the **reshape()** function, which reflects the number of replicates (three for each sample) and the total sample count. Thereafter, the code generates a dataset including the calculated means split into 125-sample chunks, representing the spectroscopic data section from 908 to 1676 nm. For each block of data, the **pinv()** function calculates the pseudo-inverse of the matrix, which is used for linear regression. Afterwards, the **block_multiply()** function multiplies this pseudo-inverse matrix by the reshaped spectroscopic data, to calculate two important correction parameters: the slope and the intercept. Finally, the original spectroscopic data are corrected by subtracting the slope and dividing by the calculated intercept. Following this step, the normalisation phase aims to minimise the influence of intrinsic variability between samples caused by unwanted influences. Such calculation is obtained first by subtracting the mean of the MSC values, and then by dividing by the standard deviation. Figure 5 shows the normalised graphs achieved from the two types of samples.

**Figure 5.**
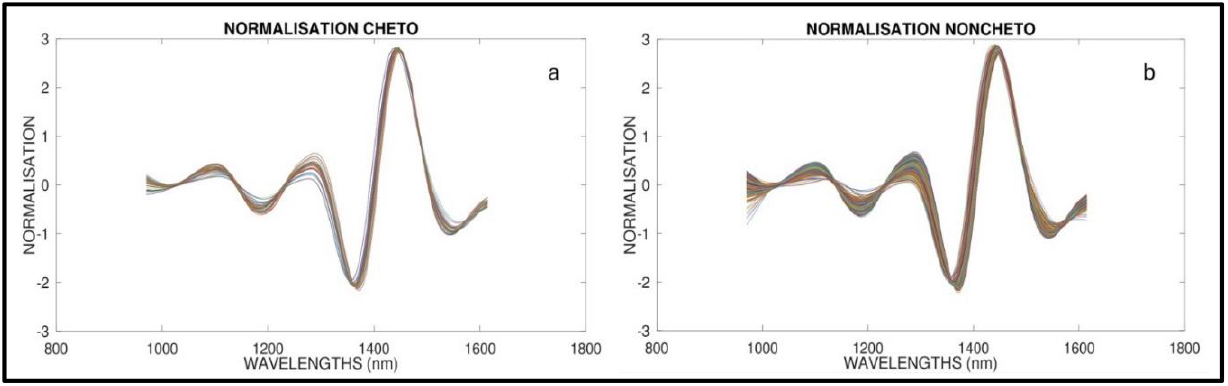
Normalised spectra of milk samples classified as “Ketotic” (a) and “Not-Ketotic” (b).

The aquagram construction and subsequent chemometric calculations involved exclusively Farm 1 (DB) and Farm 2 (BR), since no samples with a fat-protein ratio greater than 1.4 resulted from the third farm. To better understand which water absorption bands are fundamental for distinguishing milk samples, a linear graphic representation was realised at the end of this first spectral preprocessing phase, both on the total of the analysed milk samples as well as on each individual farm. This plot shows the wavelengths selected based on the most significant differences in normalised absorbance obtained by comparing the mean values between ketotic and non-ketotic sample groups. Such procedure was carried out over a restricted range of wavelengths, between 1310 and 1589 nm, a specific spectral range defined by the Water Matrix Coordinates (WAMACS), for the identification of key spectral regions linked to specific molecular conformations of water (Tsenkova, 2009). Such spectral regions reveal which water conformations are involved with the onset of ketosis, highlighting the specific interactions between water molecules and milk components (Figure 6).

**Figure 6.**
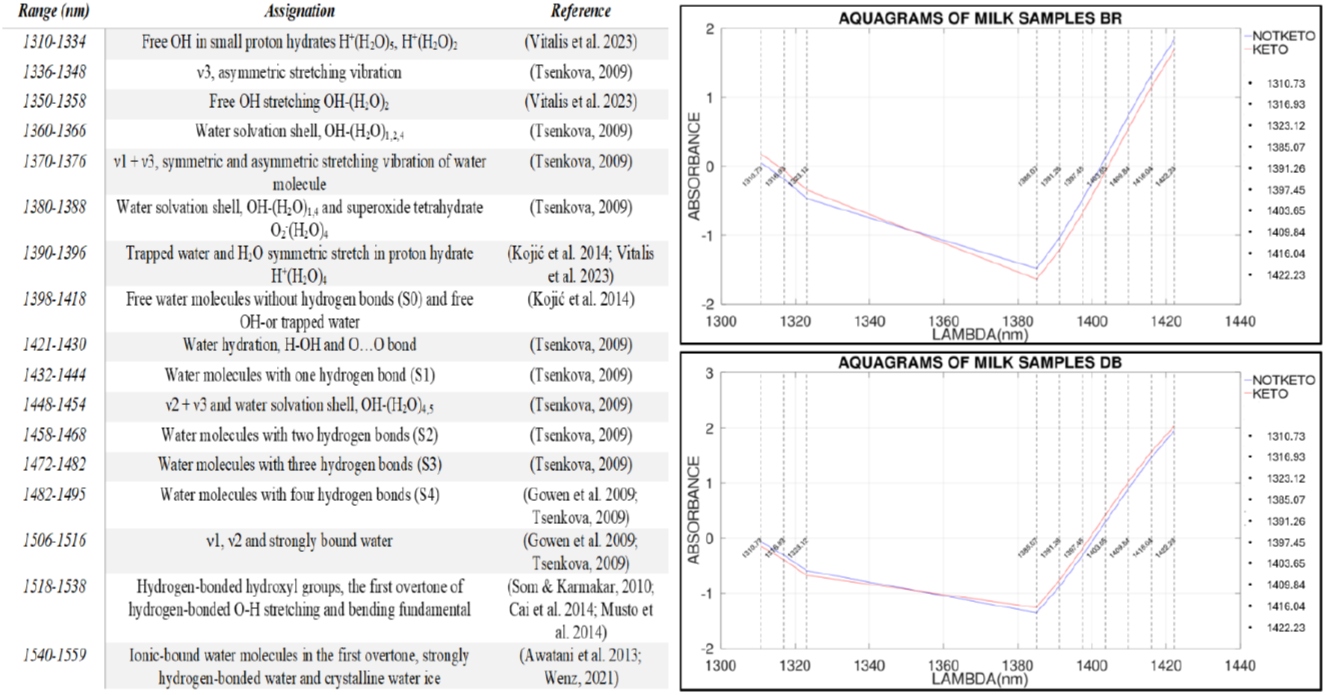
Coordinates of the molecular matrix of water with their spectroscopic and structural characterisation. On the right the aquagrams showing the normalised water absorption bands relevant for the discrimination of milk samples. The analysis includes samples from Farm 2 (BR) (top graph), samples from Farm 1 (DB) (bottom graph).

### 2. Chemometric Approach

The second section of the script applies a chemometric approach to determine a predictive model based on the water’s spectroscopic features. This passage is fundamental for a deeper comprehension of the spectral regions with the biggest role in distinguishing among the two sample categories. It is important to emphasise that all the operations executed in these passages are performed both on the total set of samples and separately for each of the farms analysed with at least one ketotic sample (in this case, only Farm 1 DB and Farm 2 BR). For this reason, the script can be adapted by adding or deleting repeated sections, depending on the distinct experimental layout.

The MSC values included in the spectral range defined by the WAMACS are thereafter concatenated with the compositional data of the analysed milk samples, contained in the text file Composition.txt, and assigned to the UnionePCA matrix. The chemometric analysis begins with the execution of the PCA by reducing the data dimensionality maintaining a small number of principal components associated with high spectral variance. This step is fundamental, both to observe a possible structure in sample cloud, and to the construction of the LDA and QDA classification models.

The principal component analysis is performed exclusively on the portion of UnionePCA containing the normalised spectral data by calling the **pca()** function. The principal components, the loadings (i.e., the contribution of each original variable on each principal component), the scores (i.e., the coordinates of the samples in the new principal component space) and the variance explained by each principal component were thus calculated. The post-processing of the PCA results calculates, for each wavelength, the sum of the loadings occurring in each principal component to determine the cumulative effect of each water wavelength absorbance on the differentiation between the ketotic and healthy groups. Such a value is important in highlighting the wavelengths whose absorbances discriminate among ketotic and healthy samples the most. The ten most significative wavelengths are then selected and saved. Finally, the obtained results are presented through 3D scatter plots (to visualise the spectra as scores) and bar plots (to visualise the loadings of the selected wavelengths on the first three principal components) (Figure 7).

**Figure 7.**
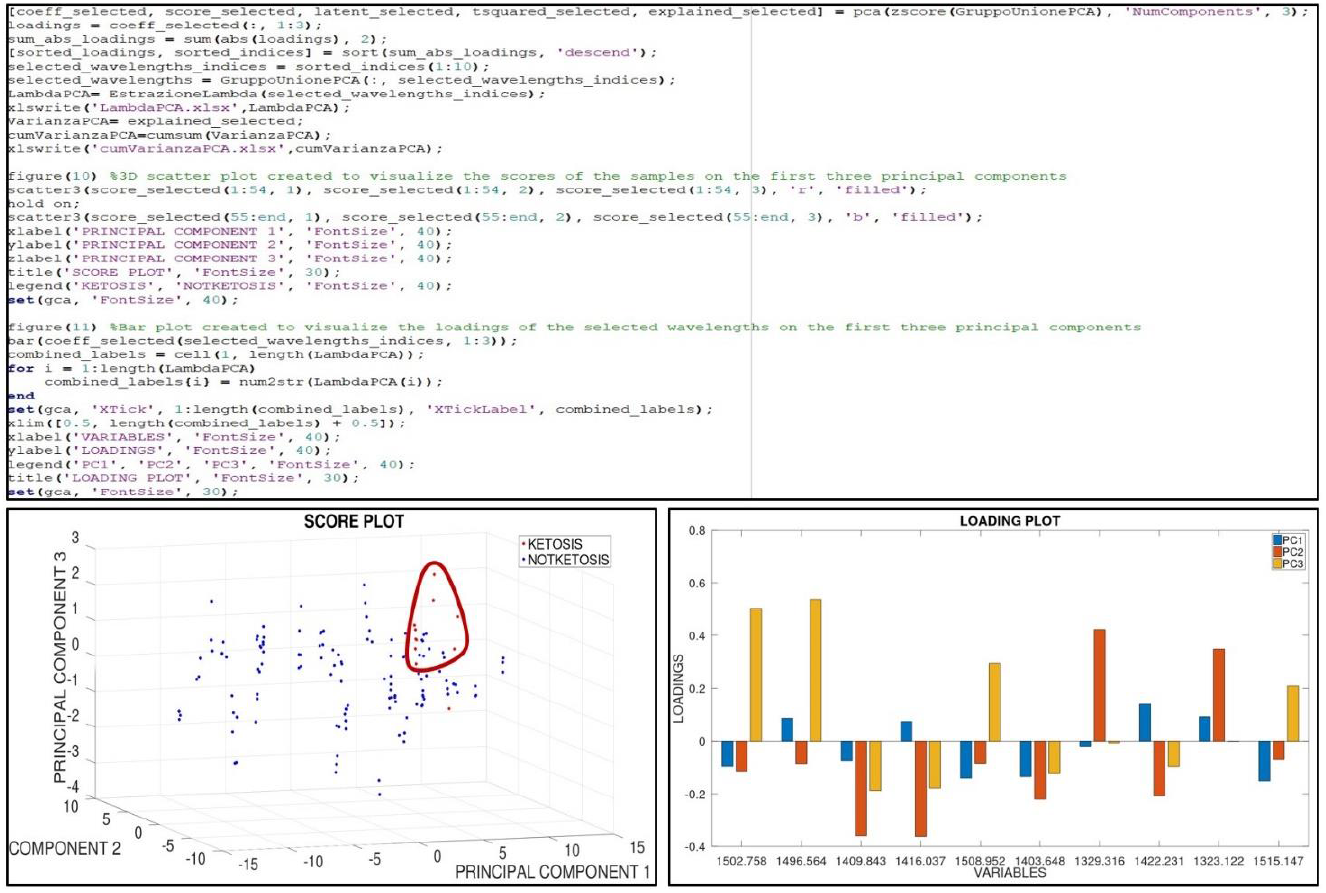
Section of the code used to execute the Principal Component Analysis. On the left, the score plot with red dots for animals with ketosis conditions in which blue dots are for healthy cattle. The loading plot with the ten most significative wavelengths selected from PCA is on the right. Both graphs belong to samples collected from Farm 2 (BR).

At this point, as mentioned, it follows the realisation of a classification model based both on the chemical parameters and also on the wavelengths picked out by PCA. This is possible thanks to the utilisation of two different classification techniques: linear and quadratic analysis.

LDA is employed to identify a linear combination of features that most efficiently distinguish between two or more classes. This technique projects the data onto a space where a straight line is used to separate the classes. Conversely, QDA is employed to identify a quadratic combination of features that most efficiently distinguishes between two or more classes. Consequently, this method endeavours to project the data onto a space where the classes can be separated by a curve, such as a parabola. Since QDA uses quadratic functions for class separation, it models curved decision boundaries, and this is useful when data are not linearly separable.

The initial part of the code performs LDA to maximise the distinction between the two categories of samples and determine the linear discriminant functions for classification. Through three nested for statement loops, the code examines all possible combinations of three wavelengths (of the 10 most significant ones extracted from the PCA), as well as the lactation number of the analysed cattle, to identify the combination that maximises the proportion of correct classifications. Specifically, for each combination of 3 wavelengths, the code determines the discriminant vector that optimises such separation. Data are thereafter projected onto the discriminant vector and normalised, while a classification threshold is calculated based on the means of the projected data. Such threshold is defined as the midpoint between the means of the projected data of the two classes. The samples are subsequently classified into the two classes by comparing the projected value with the threshold. In this way, it is possible to determine the proportion of correctly classified samples for each combination of wavelengths. Once the best combination is identified, the code displays the proportions of correctly classified samples and the linear discriminant function equation for each of the two classes, as in the example below, where the equations for Farm 1 (DB) are reported.

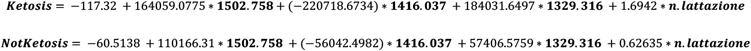

The random division of milk samples into Calibration and Validation sets plays a key role in the following classification and regression analyses. Specifically, the code randomly selects milk samples so that the calibration set comprises 60% of the ketotic samples and 60% of the not-ketotic ones; meanwhile, the validation set includes the remaining 40% belonging to the two metabolic categories. This step is performed starting from the sample indices contained in the UnionePCA matrix. Such subdivision aims to develop a robust model that could produce accurate predictions on new spectroscopic data introduced into the code.

Like LDA, the QDA classification technique examines all the hypothetical combinations of three wavelengths (from the 10 most important ones identified by PCA) as well as the lactation number of the analysed cattle, to determine the combination that maximises the proportion of correct classifications. Also with the QDA, the code calculates the mean and the covariance matrix for each class. At this point, for each sample, the script calculates a quadratic discriminant function separately for each class and stores the results in the variable “discriminant_values(c)”. This operation utilises the Mahalanobis distance, a measure that indicates how far a sample is from the mean of each class, assigning it to the class with the highest discriminant value. Once the correct classification proportion is calculated for each combination, the code saves the five best ones identified and their respective accuracy in an Excel file.

The model constructed during calibration is now applied to the validation set. It utilises the optimal combination of wavelengths (together with the cow’s lactation number) that the script picked out. The classification procedure for the validation set follows the same scheme used in the calibration. Subsequently, the algorithm generates a confusion matrix for both sets, offering an intuitive representation of the statistical classification accuracy of the model. More specifically, Figure 8 reports the number of correct and incorrect predictions realised by the model, organised into four categories: True Positives, True Negatives, False Positives and False Negatives.

**Figure 8.**
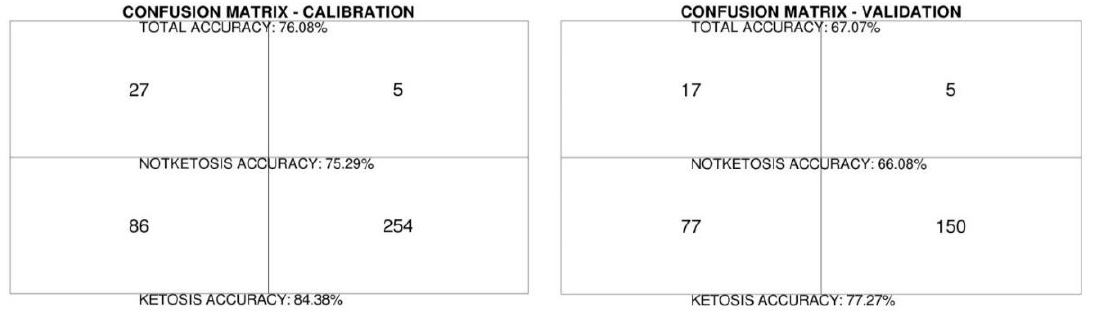
Confusion matrix obtained from the calibration set (on the left) and from the validation set (on the right) using the best combination of wavelengths extracted from the QDA model.

The last section of the code performs a PLSR, starting from the same partitioning of the calibration and validation dataset. The aim is to define a predictive model capable of estimating a target variable (fat/protein ratio - 4th column of the matrix), by utilising as predictors all the wavelengths within the WAMACS interval (6th to end columns of the matrix). To estimate the accuracy degree of the developed regression model, the code determines some performance metrics, such as the coefficient of determination (R^2^), root mean square error (RMSE), per cent bias (PBIAS), root mean square residual (RSR) and correlation index (r), on both the calibration and validation datasets. During the model calibration analysis, it is fundamental to determine the unique values of the target variable that need to be estimated, as well as the corresponding positional indices within the Calibration_Group matrix. This is made thanks to the combination of two key functions: **unique()** and **setdiff()**. The resulting duplicate values are not removed, nevertheless they are transferred to the validation dataset, a necessary step to prevent data redundancy from affecting the model performance. Once the absorption data and the target variable has been centered (by subtracting the average value from them), the PLSR can be applied by utilising the **plsregress()** function. This function, in particular, returns both the model coefficients (Coeff_cv) and the centered predicted values (Fitted_cv_centered). At this point, thanks to the utilisation of a for statement loop, the script determines the optimal model’s metrics. The resulting equation of the PLSR model, obtained by considering each predictive variable’s contribution to the fat-protein ratio calculation, is saved too. Thereafter, the model is graphically visualised with a scatter plot, a graph reporting the observed value of the target variable on the x-axis and the predicted one on the y-axis (Figure 9), with a trend line crossing the origin of the axis that helps the comparison of the results.

**Figure 9.**
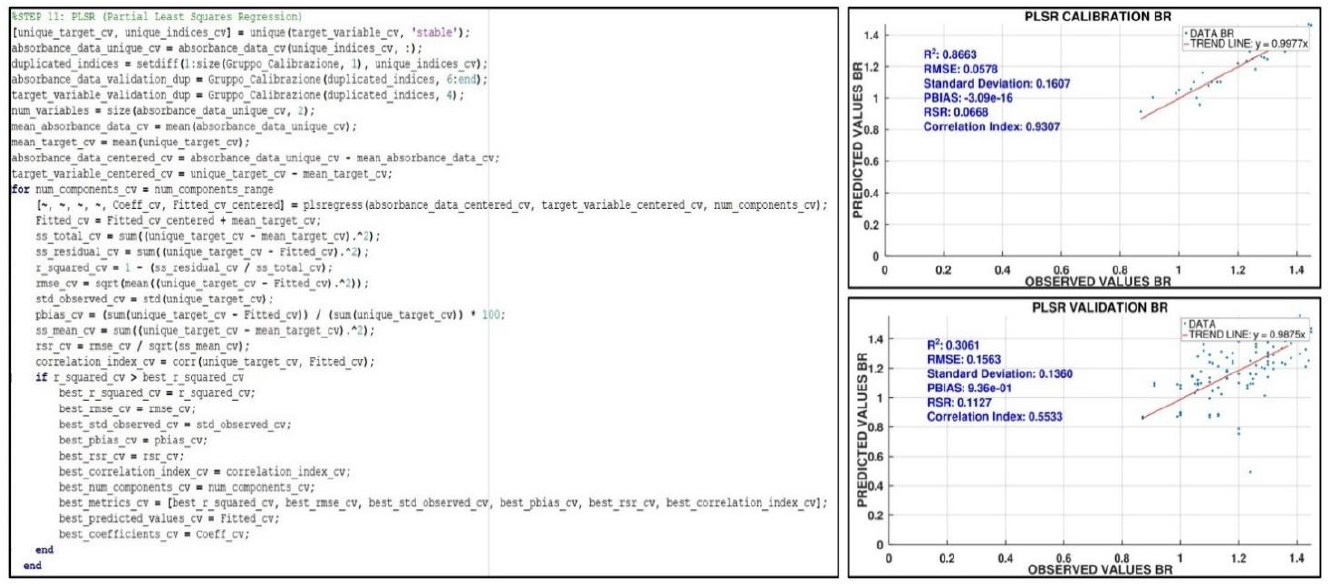
Section of the code used for the execution of the Partial Least Square Regression. On the right, two examples of scatter plot obtained from Farm 2 (BR): one for the calibration set and the other for the validation set. Each plot includes a textbox annotation that displays the ‘model’s performance metrics and the trend line.

The beta coefficients of the best calibration model are therefore employed to make predictions on the validation dataset and estimate the target fat-protein ratio variable. Similarly to the calibration, during the model validation phase the script calculates the model performance metrics. Moreover, the code realised determines the normalised and also the loading weights derived from the PLSR model’s coefficients. Such phase aims to emphasise the contribution of each water absorption band to the model’s predictive performance. In particular, their role during the onset of the disease is further investigated by analysing the overall contribution from each individual farm. Specifically, the previously calculated normalised weights are thereafter summed, and the five most significant wavelengths in predicting the target variable are displayed in a bar chart (Figure 10).

**Figure 10.**
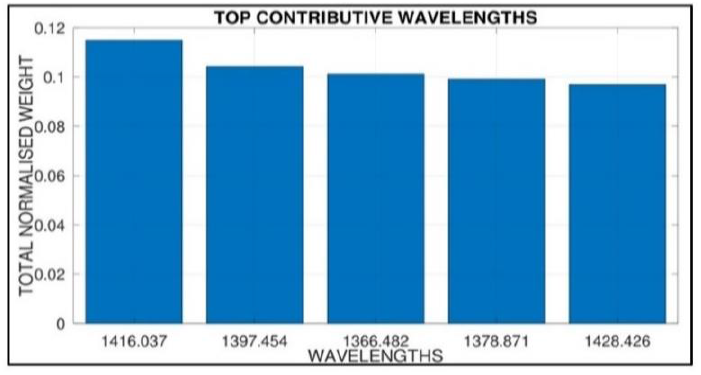
Bar plot displaying the top five total weights, obtained by summing the contributions of all farms to verify the wavelengths mainly involved in the observed variations of the target variable.

## Conclusions

Spectral preprocessing and chemometric techniques allow the identification of all the bands that could be used to describe changes in water’s spectral pattern linked to early ketosis signs. Therefore, aquaphotomics can be used to classify milk samples according to the animal’s metabolic profile. Nevertheless, this code can constitute a versatile starting point, easy to apply with minimal adjustments for different purposes of spectral analysis, adapting to any analytical application that employs aquaphotomics as an investigation method.

## Acknowledgements

The authors acknowledge the Department of Economics, Engineering, Society and Business Organization (DEIM) in cooperation with the Department of Agriculture and Forestry Sciences (DAFNE) of the University of Tuscia for this work carried out as part of the PhD in Engineering for Energy and Environment – Biosystems and Environment.

## References

1) Awatani T, Midorikawa H, Kojima N, Ye J, & Marcott C 2013 Morphology of water transport channels and hydrophobic clusters in Nafion from high spatial resolution AFM-IR spectroscopy and imaging. Electrochemistry Communications, 30, 5–8. 10.1016/j.elecom.2013.01.021

2) Cai CB, Tao YY, Wang B, Wen MQ, Yang HW, & Cheng YJ 2014 Investigating the adsorption process of isoamyl alcohol vapor onto silica gel with near-infrared process analytical technology. Spectroscopy Letters, 48, 190–197. 10.1080/00387010.2013.872668

3) Deniz A, Aksoy K, Metin M 2020 Transition period and subclinical ketosis in dairy cattle: association with milk production, metabolic and reproductive disorders and economic aspects. Medycyna Weterynaryjna, 76(09):495–502. 10.21521/mw.6427

4) Gowen AA, Tsenkova R, Esquerre C, Downey G, & O’Donnell O’Donnell CP 2009 Use of near Infrared Hyperspectral Imaging to Identify Water Matrix Co-Ordinates in Mushrooms (Agaricus Bisporus) Subjected to Mechanical Vibration. Journal of Near Infrared Spectroscopy, Vol. 17, pp. 363–371. DOI:10.1255/jnirs.860

5) Kaniyamattam K, De Vries A 2014 Agreement between milk fat, protein, and lactose observations collected from the Dairy Herd Improvement Association (DHIA) and a real-time milk analyzer. Journal of Dairy Science, 97(5):2896–908. doi: 10.3168/jds.2013-7690

6) Kojić D, Tsenkova R, Tomobe K, Yasuoka K, & Yasui M 2014 Water confined in the local field of ions. ChemPhysChem 15 (18):4077–4086. doi: 10.1002/cphc.201402381

7) Kovacs Z, Muncan J, Veleva P, Oshima M, Shigeoka S, & Tsenkova R 2022 Aquaphotomics for monitoring of groundwater using short-wavelength near-infrared spectroscopy. Spectrochimica Acta - Part A: Molecular and Biomolecular Spectroscopy, 279. 10.1016/j.saa.2022.121378

8) Loiklung C, Sukon P, Thamrongyoswittayakul C 2022 Global prevalence of subclinical ketosis in dairy cows: A systematic review and meta-analysis. Research in Veterinary Science, 144, 66–76. 10.1016/j.rvsc.2022.01.003

9) Luck WA 1973 Structure of Water and Aqueous Solutions. Proceedings of the International Symposium Marburg, 248–284. 10.1002/bbpc.19760800719

10) Muncan J, & Tsenkova R 2019 Aquaphotomics-From Innovative Knowledge to Integrative Platform in Science and Technology. Molecules 24(15):2742. 10.3390/molecules24152742

11) Musto P, Galizia M, Pannico M, Scherillo G, & Mensitieri G 2014 Time-resolved Fourier transform infrared spectroscopy, gravimetry, and thermodynamic modeling for a molecular level description of water sorption in poly(ε-caprolactone). The Journal of Physical Chemistry B, 118(26). DOI: 10.1021/jp502270h

12) Roger JM, Mallet A, & Marini F 2022 Preprocessing NIR Spectra for Aquaphotomics. Molecules (Vol. 27, Issue 20). MDPI. 10.3390/molecules27206795

13) Setti G 2017 Chetosi e cellule, nemiche del caseificio. Informatore Zootecnico n.12/2017

14) Som T, & Karmakar B 2010 Structure and properties of low-phonon antimony glasses and nano glass-ceramics in K2O– B2O3–Sb2O3 system. Journal of Non-Crystalline Solids, 356:987–99. doi: 10.1016/J.JNONCRYSOL.2010.01.026

15) Todeschini R 1998 Introduzione alla chemiometria. Napoli: Edises; EAN: 9788879591461, ISBN: 8879591460

16) Tsenkova R 2009 Introduction aquaphotomics: Dynamic spectroscopy of aqueous and biological systems describes peculiarities of water. Journal of Near Infrared Spectroscopy (Vol. 17, Issue 6, pp. 303–314). 10.1255/jnirs.869

17) Tsenkova R, & Muncan JS 2021 Aquaphotomics for Bio-diagnostics in Dairy. Applications of Near-Infrared Spectroscopy. Springer Nature Singapore, ISBN: 978-981-16-7114-2

18) Tsenkova R, Muncan JS, Pollner B, & Kovacs Z 2018 Essentials of aquaphotomics and its chemometrics approaches. Frontiers in Chemistry, 6(AUG). 10.3389/fchem.2018.00363

19) Van de Kraats EB, Munćan JS, & Tsenkova R 2019 Aquaphotomics Origin, concept, applications and future perspectives. An International Journal of the History of Chemistry, 3(2), 13–28. 10.13128/Substantia-702

20) Vitalis F, Muncan J, Anantawittayanon S, Kovacs Z, & Tsenkova R 2023 Aquaphotomics Monitoring of Lettuce Freshness during Cold Storage. Foods, 12(2). 10.3390/foods12020258

21) Wenz JJ 2021 Influence of steroids on hydrogen bonds in membranes assessed by near infrared spectroscopy. Biochimica et Biophysica Acta (BBA) – Biomembranes, 1863, 183553. 10.1016/j.bbamem.2021.183553

