## Supplementary Table S1 for "Identification of ketotic cattle based on milk spectroscopic characteristics: a chemometric modelling approach built upon the principles of Aquaphotomics"

### SUPPLEMENTARY FILE

***Supplementary Table S1*** *Overview of all the functions used in the spectroscopic data processing code categorised according to their function.*

#### *Mathematical and Statistical Functions related to Matrix and Vector Calculations*

***abs****: Calculates the absolute value of elements*
***ceil****: Rounds elements up to the nearest integer*
***corr****: Calculates the correlation between two variables*
***cov****: Calculates the covariance matrix*
***det****: Calculates the determinant of a matrix*
***eig****: Calculates eigenvalues and eigenvectors of a matrix*
***inv****: Calculates the inverse of a matrix*
***log****: Calculates the natural logarithm of elements*
***max****: Finds the maximum value in an array*
***mean****: Calculates the mean of elements*
***min****: Finds the minimum value in an array*
***norm****: Calculates the norm of a vector or matrix*
***pca****: Performs Principal Component Analysis*
***pinv****: Calculates the pseudo-inverse of a matrix*
***plsregress****: Performs Partial Least Squares Regression*
***sgolayfilt****: Applies Savitzky-Golay filtering to data*
***sort****: Sorts elements in an array*
***sqrt****: Calculates the square root of elements*
***std****: Calculates the standard deviation of elements*
***sum****: Calculates the sum of elements*
***zscore****: Calculates the Z-score of elements*

#### *Input/Output Functions*

***disp****: Displays information in the command window*
***fprintf****: Prints formatted output to screen or file*
***load****: Loads data from a file*
***num2cell****: Converts numbers to cells*
***num2str****: Converts numbers to strings*
***sprintf****: Formats data into strings*
***write_excel_output****: Custom function to write data to Excel*

***xlsfinfo**: Provides information about Excel files*
***xlsread**: Reads data from an Excel file*
***xlswrite**: Writes data to an Excel file*

### *Visualisation and Graphics Functions*

***annotation**: Adds annotations to figures*
***bar**: Creates a bar chart*
***caxis**: Sets the colour axis limits for colour maps*
***colormap**: Defines the colour map for figures*
***figure**: Creates a new figure window*
***get**: Retrieves properties of graphic objects*
***grid on**: Turns on the grid in figures*
***hold off**: Disables holding the current graph for additional plotting*
***hold on**: Keeps the current graph for additional plotting*
***imagesc**: Displays a matrix as a scaled image*
***legend**: Adds a legend to figures*
***meshgrid**: Creates a grid of coordinates for 3D plotting*
***plot**: Creates a 2D plot*
***rectangle**: Adds rectangles to figures*
***scatter**: Creates a scatter plot*
***set**: Sets properties of graphic objects*
***text**: Adds text to figures*
***title**: Adds a title to figures*
***xlabel**: Adds a label to the x-axis*
***xlim**: Sets the limits of the x-axis*
***xticklabels**: Defines tick labels on the x-axis*
***xticks**: Defines ticks on the x-axis*
***ylabel**: Adds a label to the y-axis*
***ylim**: Sets the limits of the y-axis*

### *Data Manipulation and Transformation Functions*

***block_multiply**: Multiplies matrices in blocks*
***cat**: Concatenates arrays along a specified dimension*
***cell**: Creates cell arrays*
***diag**: Extracts or creates a diagonal matrix*
***intersect**: Finds the intersection between two arrays*
***ismember**: Checks if elements are members of an array*
***length**: Returns the maximum dimension of an array*
***ones**: Creates an array where all elements are set to 1*
***permute**: Permutes the dimensions of an array*
***reshape**: Reshapes an array*
***setdiff**: Finds the difference between two arrays*
***size**: Returns the dimensions of an array*
***strcat**: Concatenates strings*
***unique**: Finds unique values in an array*

### *Initial and Package Interface Functions*

***clear all**: Clears all variables in the workspace*
***pkg load**: Loads packages in Octave*
